# Divergent associations between auditory activation and inhibition task performance in children and adults

**DOI:** 10.1101/2022.12.16.520726

**Authors:** Sam van Bijnen, Tiina Parviainen

**Author notes:** Corresponding author at, Department of Psychology, University of Jyväskylä, Kärki, Mattilanniemi 6, FI-40014 Jyväskylän yliopisto, Finland. Declarations of interest: none.

## Abstract

Adults and children show remarkable differences in cortical auditory activation which, in children, have shown relevance for cognitive performance, specifically inhibitory control. However, it has not been tested whether these differences translate to functional differences in response inhibition between adults and children. We recorded auditory responses of adults and school-aged children (6-14y) using combined magneto- and electroencephalography (M/EEG) during passive listening conditions and an auditory Go/No-go task. The associations between auditory cortical responses and inhibition performance measures diverge between adults and children; while in children the brain-behavior associations are not significant, or stronger responses are beneficial, adults show negative associations between auditory cortical responses and inhibitory performance. Furthermore, we found qualitative differences in auditory responses between adults and children; the late (∼200 ms post stimulation) adult peak activation shifts from auditory to frontomedial areas. In contrast, children show prolonged obligatory responses in the auditory cortex. Together this likely translates to a functional difference between adults and children in the cortical resources for performance consistency in auditory-based cognitive tasks.

## Introduction

Research dedicated to understanding the development of the central auditory system was crucial to reveal the important cortical timings in auditory perception in adults and children. It led to the current understanding that cortical auditory evoked responses change substantially from childhood to adulthood (Paetau et al., 1995; Johnstone et al., 1996; Ponton et al., 2000; Ponton et al., 2002; Wunderlich and Cone-Wesson, 2006). These changes coincide with grey and white matter transitions (Moore & Linthicum Jr, 2007) that interact with synaptic signaling to affect the timing and amplitude during development. Indeed, the evoked responses can be used as a measure of cortical network efficiency, as they rely on temporal accuracy of the cortex to respond to stimuli. Likewise, the timings of the activation pattern are an indicator of (auditory) development (Parviainen et al., 2011; Hämäläinen et al., 2013; van Bijnen et al., 2019). The characteristics of these auditory evoked responses have important links with behavioral skills both in adults and in children (e.g., Näätänen, 1990; Jonhstone et al., 1996; Parviainen et al., 2011; van Bijnen et al., 2022), which makes them particularly useful in understanding incremental cognitive competency in human development.

The time-windows of neural activation after auditory stimulation are, however, remarkably different in children and adults. The polyphasic adult waveform, as measured by electro- and magnetoencephalography (EEG/MEG), is characterized by intermittent positive deflections at ∼50ms (P1) and ∼150ms (P2) and a prominent negative deflection at ∼100ms (N1). In contrast, children typically show a biphasic response pattern consisting of a positive deflection at ∼100ms (P1) and a prolonged negative deflection at ∼250ms (N250) (Picton et al., 1974; Ponton et al., 2000; Albrecht et al., 2000; Čeponienė et al., 2002; Takeshita et al., 2002; Wunderlich et al., 2006; Sussman et al., 2008; Orekhova et al., 2013; Ruhnau et al., 2011; Yoshimura et al., 2014). In the early time-window, the typical adult N1-P2 complex emerges during adolescence (Sussman et al., 2008). However, the activity in the later time-window (∼250ms) attenuates and is specific to the child-brain (Sussman et al., 2008; Parviainen et al., 2011; Parviainen et al. 2019).

We have previously shown that the later time-window (∼250ms) of auditory activation reflects behaviorally meaningful processes, as it contributes to variance in response inhibition in children even when measured during passive listening conditions (van Bijnen et al., 2022). While there seems to be converging evidence on the behavioral relevance of the later time-window for both auditory-based and more general skills during childhood (van Bijnen et al., 2022; Johnstone et al., 1996), the role of the early time-window responses remains unclear. From one perspective, the early responses (< 200ms) are exogenous (obligatory) activity and evoked in both adults and children, without any task or attention. However, there are indications that neural processing in this time-window contributes to language and communication ability (Parviainen et al., 2005; Yoshimura et al., 2014) and arousal or attention regulation (Orekhova et al., 2012) in children.

We have suggested that children and adults employ divergent brain mechanisms for a consistent performance in auditory based cognitive tasks. This claim was based on the finding that children relied strongly on the child-specific auditory activation at 250ms for a consistent performance on (auditory) inhibition tasks (van Bijnen et al., 2022). In contrast, it is well known that adults rely on frontal/medial regions of the cerebral cortex during this time-window for inhibitory and cognitive control processes (Nieuwenhuis et al., 2003; Huster et al., 2010; Falkenstein et al., 1999; Smith et al., 2007; Botvinick et al., 2004; Chambers et al., 2009). Yet, as far as we know, no studies have addressed this discrepancy.

Qualitative differences in neural activity pattern complicates comparisons between children and adults in M/EEG studies. Combing M/EEG is favorable because MEG has better signal-to-noise ratio, while EEG provides a better account of deeper and radial sources (Piastra et al., 2020; Baillet, 2017). Therefore, even within adults the different deflections are best picked up by either MEG or EEG depending on the source locations and orientations (Shahin et al., 2007; Piastra et al., 2020). Indeed, MEG auditory source waveforms vary across individuals depending on the Heschl’s gyrus gyrification type (Benner et al., 2017) and the anatomical organization of the auditory cortex (Shaw et al., 2013). In children there is the added difficulty of variability in the developmental stages even within a narrow age-range. Therefore, source locations and orientations might differ between individuals and combined MRI and M/EEG better accounts for these individual differences.

In the present study, we aimed to investigate possible differences in both the spatio-temporal characteristics of activation and the brain-behavior associations between children and adults in the early auditory activation supporting response inhibition. We focused on the early auditory responses that are prevalent in both children and adults and contrasted an auditory Go/No-go task with a passive listening task (Fig 1.).

**Figure 1.**
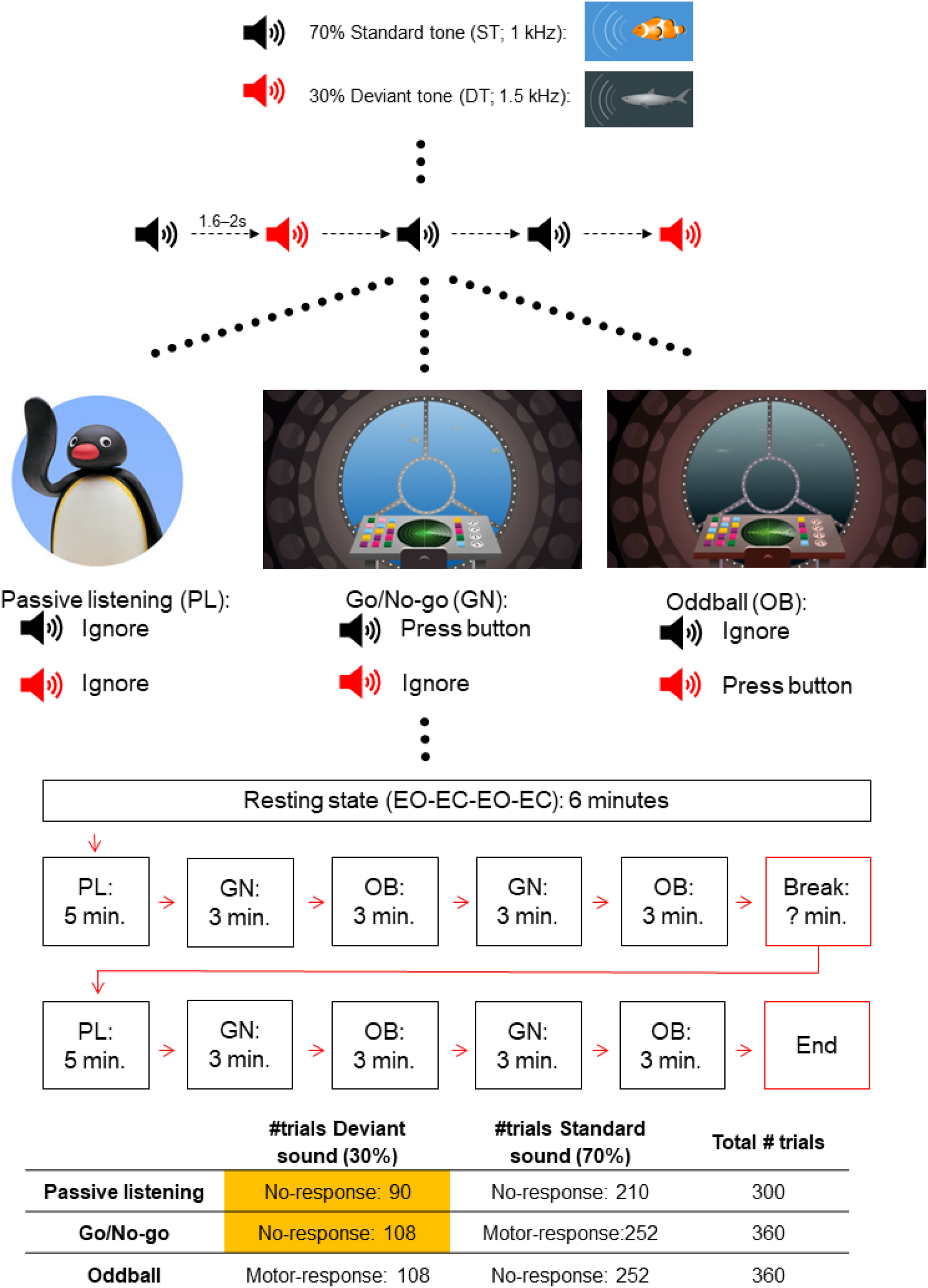
Experimental design and procedure. In this study, we specifically focused on the passive listening (PL) Go/No-go (GN) comparison (marked in yellow) to limit the number of statistical tests.

## Materials and Methods

### Participants

Seventy-eight children that were recruited through schools and the National registry and sixteen adults participated in this study. Eleven children and zero adults were excluded due to data quality issues, not finishing the experiment, or structural abnormalities in the MRI. The final dataset consisted of sixty-seven children (36 boys) aged between 6-14 years (M = 10.2, SD = 1.4). and sixteen adults (13 women) aged between 20-30 years (M = 24.8, SD = 3.4). All participants had normal hearing as tested with an audiometer. None of the participants had neurological disorders or were on medication affecting the central nervous system. The study was approved by the Ethics Committee of the University of Jyväskylä. An informed consent was obtained from all children and their parents, and the adults in accordance with the Declaration of Helsinki. All participants received compensation for participation (movie ticket or gift card).

The children (but not adults) in this study have previously been described in van Bijnen et al. (2022, which focused on the child-specific N250 activation. Here, we focused on the transient (early) auditory responses that are typically present in both the adults and children.

### Tasks and procedure

The task was presented as a game where the participants were asked to help science by studying and protecting the clownfish population. Studio Dennis Parren (www.dennisparren.com) created a visual environment (video) resembling a submarine with a captain giving instructions. All stimuli were controlled by PsychoPy (V3.2) (Peirce et al., 2019) running on a Linux desktop PC. Auditory stimuli were delivered binaurally to the subject through plastic tubes and earpieces using an MEG-compatible hi-fidelity sound system.

First, we measured resting state activity using two alternating 1.5 minutes eyes open and closed sessions. Subsequently, the task started with the first passive listening (PL) oddball task. Participants listened to a continuous stream of deviant (DT; 1.5 kHz) and standard (ST; 1 kHz) tones with a 70‒ms duration (10‒ms rise/fall time) separated with an inter-stimulus interval varying between 1.6 and 2.0‒s and were instructed to ignore both tones while they could watch a silent stop-motion video (“Pingu”). After the first PL task participants completed two blocks of alternating Go/No-go (GN) and oddball detection (OB) tasks before the break. Stimuli were identical to the PL task, but participants were asked to either respond to the deviant tone (OB) or the standard tone (GN). After a break, participants completed the same blocks of the PL, GN & OB tasks as before the break. The complete procedure is shown in figure 1.

### Behavioral assessment

Cognitive skills were tested on a separate visit. The behavioral tests included subtests of Wechsler Intelligence Scales for Children Third edition (Wechsler, 1999) or Wechsler Adult Intelligence Scale, which was used to evaluate the general level of cognitive skills, and the Stop Signal Task (SST) from the Cambridge Neuropsychological Automated Test Battery (CANTAB).

Of the Wechsler Intelligence scale, the similarities and block design subtests were used to evaluate the general level of verbal and nonverbal reasoning, respectively, and digit span (backward/forward) was used as a measure verbal short-term memory. The coding test and the symbol search task were used to evaluate in general the speed of processing, visuomotor coordination, and attention.

The SST was used to quantify a stop-signal reaction time (SSRT) in adults and children. The SSRT is a behavioral performance measure of response inhibition, or the ability to stop an already initiated motor response which we correlated with the auditory brain activation. In the SST, the participant must respond to an arrow stimulus by selecting one of two options depending on the direction in which the arrow points. The test consists of two parts: in the first part, the participant is first introduced to the test and told to press the left-hand button when they see a left-pointing arrow and the right-hand button when they see a right-pointing arrow. There is one block of 16 trials for the participant to practice this. In the second part, the participant is told to continue pressing the buttons when they see the arrows, but if they hear an auditory signal (a beep), they should withhold their response and not press the button. The task uses a staircase design for the stop signal delay (SSD), allowing the task to adapt to the performance of the participant, narrowing in on the 50% success rate for inhibition. The SST quantifies a stop-signal reaction time (SSRT);

### M/EEG and MRI data analysis

Brain responses were recorded using a 306-channel MEG system and the integrated EEG system (Elekta Neuromag® TRIUX™, MEGIN Oy, Helsinki, Finland). M/EEG data were sampled at 1000Hz and filtered at 0.1–330 Hz. Simultaneous 32-channel EEG and vertical and horizontal electrooculograms (EOG) were recorded with an online reference on the right earlobe. A head position indicator (HPI) continuously monitored the head position in relation to the MEG sensors using five HPI coils, three anatomical landmarks (nasion, left and right preauricular points) and 150 distributed scalp points.

MEG data were first processed and converted to the mean head position with the temporal signal space separation (tSSS) and movement compensation options, implemented in the MaxFilter™ program (version 3.0; MEGIN Oy, Helsinki, Finland). Using MNE-python (Gramfort et al., 2013; Gramfort et al., 2014) data were low-pass filtered, re-referenced to the average (EEG) and bad channels and data segments were excluded. Epochs were created from –0.2 to 0.8 s relative to stimulus onset and a baseline correction was applied. Epochs from incorrect responses during the task and large MEG signals (> 4 pT/cm for gradiometers, > 5 pT for magnetometers) were rejected. Independent components representing ocular and/or cardiac artifacts were suppressed with ICA (Hyvärinen and Oja, 2000). Epochs were checked manually and with *autoreject (EEG)* (Jas et al., 2017) to repair or exclude bad epochs.

We used cortically-constrained, depth-weighted (*p* = 0.8) L2 minimum norm estimate (Hämäläinen and Ilmoniemi, 1994) with a loose orientation constraint (0.2) to characterize the source currents. T1- and T2-weighted 3D spin-echo MRI images were collected with a 1.5 T scanner (GoldSeal Signa HDxt, General Electric, Milwaukee, WI, USA) using a standard head coil and with the following parameters: TR/TE = 540/10 ms, flip angle = 90º, matrix size = 256 x 256, slice thickness = 1.2 mm, sagittal orientation. The cortical surface was constructed from the individual MRIs with the Freesurfer software (RRID: SCR_001847, Martinos Center for Biomedical Imaging, http://freesurfer.net; Dale et al., 1999; Fischl et al., 1999; Fischl et al., 2001). Next, the M/EEG data were registered to the individual structural data with MNE coregistration using the anatomical landmarks, digitized EEG electrodes and additional scalp points. The forward solution for the source space was constructed using a three-layer BEM with the following conductivity values for brain/CSF, skull and scalp: .3, .006 and .3 for adults and .33, .0132 and .33 for children. The noise covariance matrix was calculated from the individual epochs 200-ms pre-stimulus baseline, using a cross validation method implemented in MNE. The MEG and EEG data were combined into a single inverse solution by a whitening transformation using the covariance matrix (Engemann and Gramfort, 2015).

The final source waveforms were computed as the mean value within the transverse temporal gyrus (label 30 in the Desikan-Killianty Atlas; Desikan et al., 2006). The time-window for the extraction of amplitude values were based on the grand averages (figure 3) where the time-windows for the main peaks corresponded to those reported in earlier literature in children and adults (Picton et al., 1974; Ponton et al., 2000; Albrecht et al., 2000; Orekhova et al., 2013; Yoshimura et al., 2014). In children, we extracted the maximum value between 76–104ms (P1), minimum value between 108–140ms (N1) and the difference between N1 and max value between 148–200ms (P2). In adults we extracted the same P1-N1-P2 responses in different time-windows; 40–76ms, 92–120ms and 140–200ms respectively.

### Statistical analysis

We focused on the adult and child differences in the Passive Listening (PL) vs Go/No-Go task (figure 1). Task requirements in the Go/No-Go task specifically engaged inhibition. Moreover, our earlier (van Bijnen et al., 2022) study indicated that although both oddball and go/no-go comparison both yielded behaviorally significant activation, the Go/No-go showed stronger correlations. In order to limit unnecessary statistical testing, we therefore analyzed the deviant tones of these tasks.. Importantly, between the tasks, the stimuli (DT), probability (30 percent) and motor response (None) were identical. Auditory responses (P1-N1-P2) were analyzed separately; models contained one of the brain responses (P1, N1 or P2) as dependent variable at a time. Two within-subject independent (hemisphere (left, right) and task (passive, no-go)) and one between-subject variable (group; children vs adults) were included in the model. Models were estimated by using Multigroup analysis with Mplus statistical package (Version 8.4) and using a full information maximum likelihood (FIML) estimation method with robust standard errors (MLR). All available data were used in the analyses and missing data were assumed to be Missing at Random (MAR) (Muthén & Muthén, 2012). Interactions and main effects were estimated by using additional parameters of model.

Subsequently, we used (partial) correlations (corrected for age in children) to test for relevant brain-behavior associations of the auditory responses. We included the following behavioral performance measures: intra-individual reaction times (ICV; calculated as SDRT/mean RT), response accuracy (RA; calculated as the square root of the error%) and the stop-signal reaction time (SSRT). Correlational analysis was performed with SPSS statistics 25.

## Results

### Descriptive statistics of cognitive skills and behavioral performance

Descriptive statistics of the children’s performance during the M/EEG experiment and their cognitive skills as per the behavioral assessment session are presented in Table 1. The subtests of the Wechsler Intelligence Scale were included to examine whether participants are within typical range of cognitive performance and were not associated with the brain responses of children and adults (p > .05).

**Table 1.**
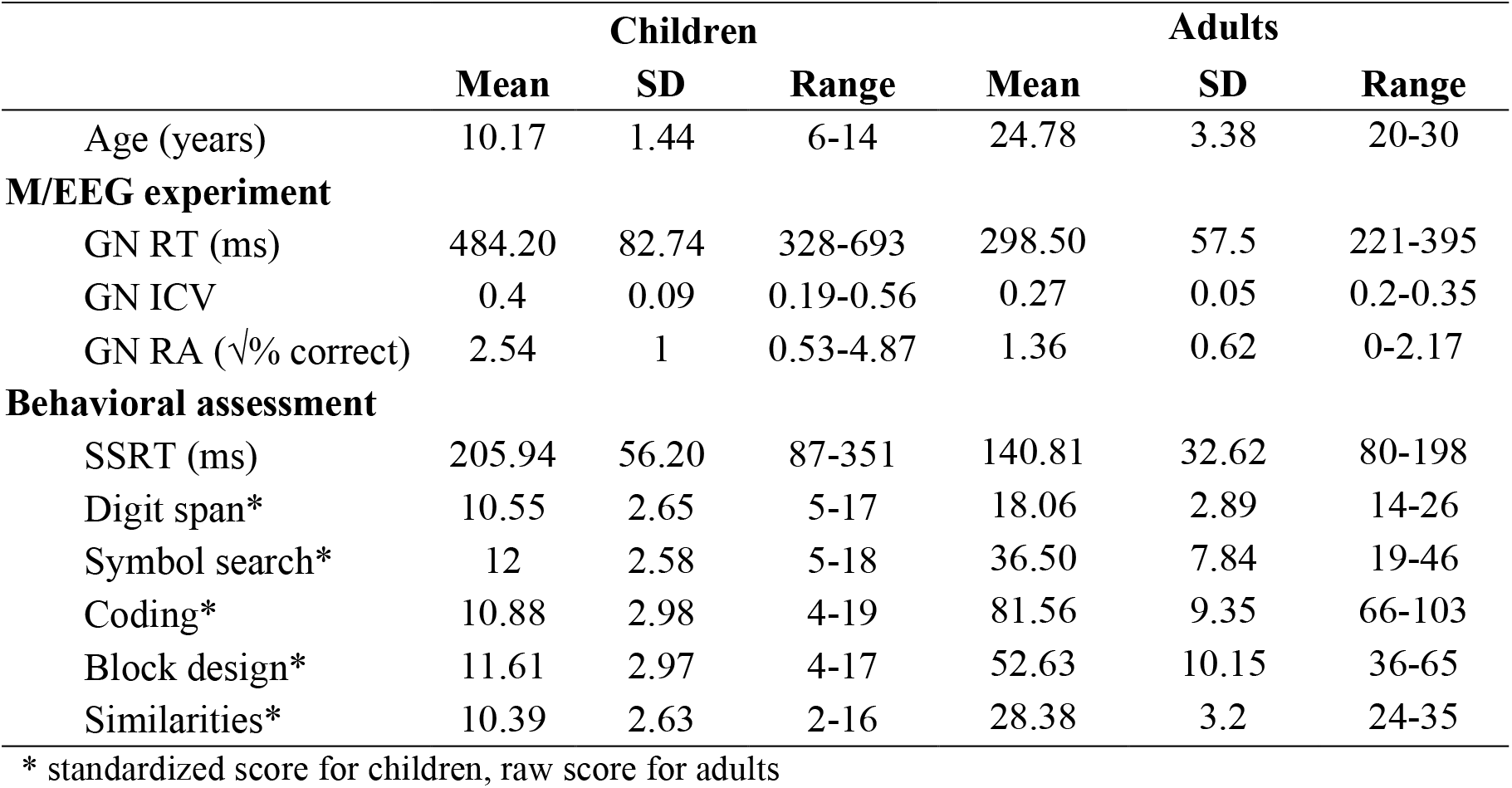
Mean, standard deviation (SD) and range of behavioral performance measures. Reaction times (RT), intra-individual coefficient of variation (ICV) and response accuracy (RA) gathered from the Go/No-go task (GN) and the Oddball task (OB). Stop-signal reaction time (SSRT) was gathered from the stop-signal task during the behavioral assessment.* standardized score for children, raw score for adults

### Age related differences in the auditory evoked responses

Figure 2 shows the measured neuromagnetic responses to the standard tones in the passive listening task at MEG sensor level (gradiometers). For visualization purposes, groups were separated by age (< 10 years old, > 10 years old and adults). The activation pattern in younger children indicate three separate peaks at ∼85ms, at ∼120ms and at ∼250ms, but with some overlap in timing especially in the youngest group. Furthermore, responses appear different in the two hemispheres in children; with the P1m predominantly showing in the left-hemisphere, the N1m predominantly showing in the right-hemisphere and the late N250m showing bilaterally. In contrast, the activation peaks in adults occur somewhat earlier, at ∼60ms and at ∼110ms, and the peak at ∼250ms is clearly diminished. Adults also show less hemispheric differences than children.

**Figure 2.**
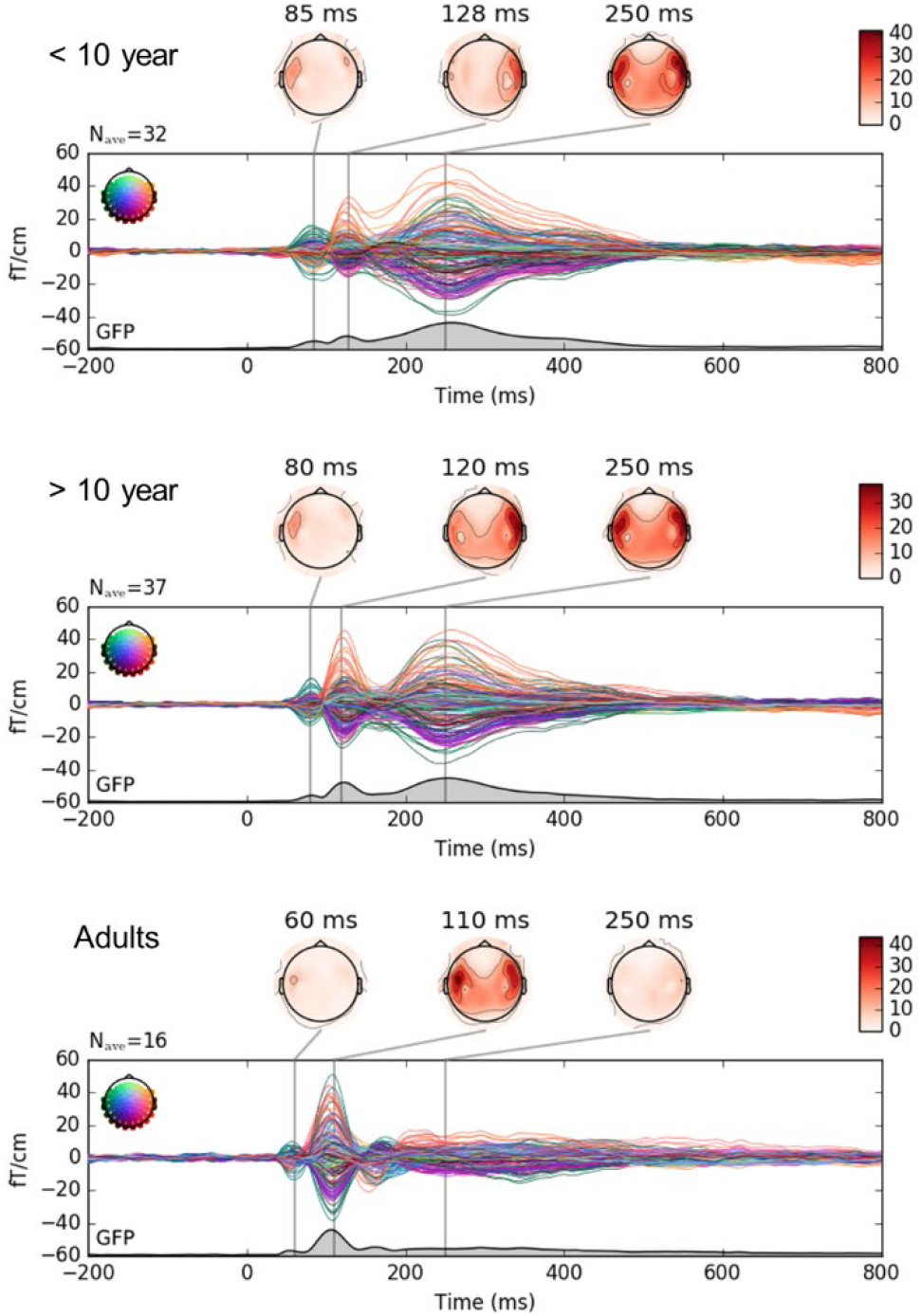
Developmental (age) differences in auditory brain responses to the passive listening (PL) standard tone (ST) as measured by the MEG gradiometers. Groups divided for illustration purposes between children younger than 10 (top), older than 10 (middle) and adults (bottom).

### Differences in child and adult auditory responses

The localization of auditory activation in children indicates that the peaks across the entire timeline of activation all reflect cortical currents in the temporal regions irrespective of task and time-window (figure 3 & 4). In contrast, in adults the early peak at 100ms reflects activation in the temporal regions and the later activation at ∼200-300ms reflects activation in the medial regions of the cerebral cortex (e.g., cingulate cortex; figure 3).

**Figure 3.**
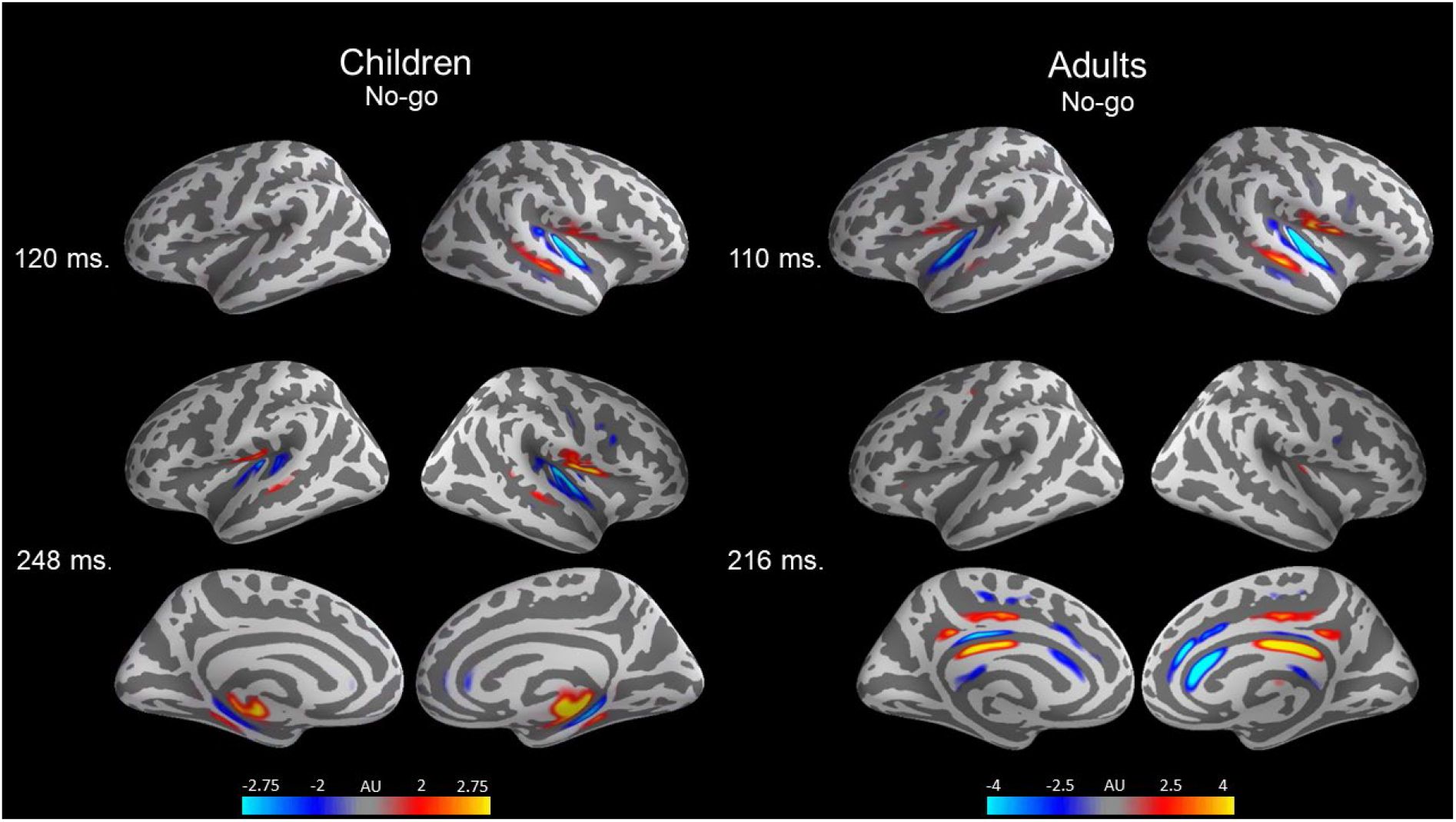
Grand average 3D visualization of the No-go (deviant tone) M/EEG combined source estimates for all children (right) and adults (left). 3D-plots are presented for the two most prominent time-windows of activation in children (120ms and 248ms) and adults (110ms and 216ms).

**Figure 4.**
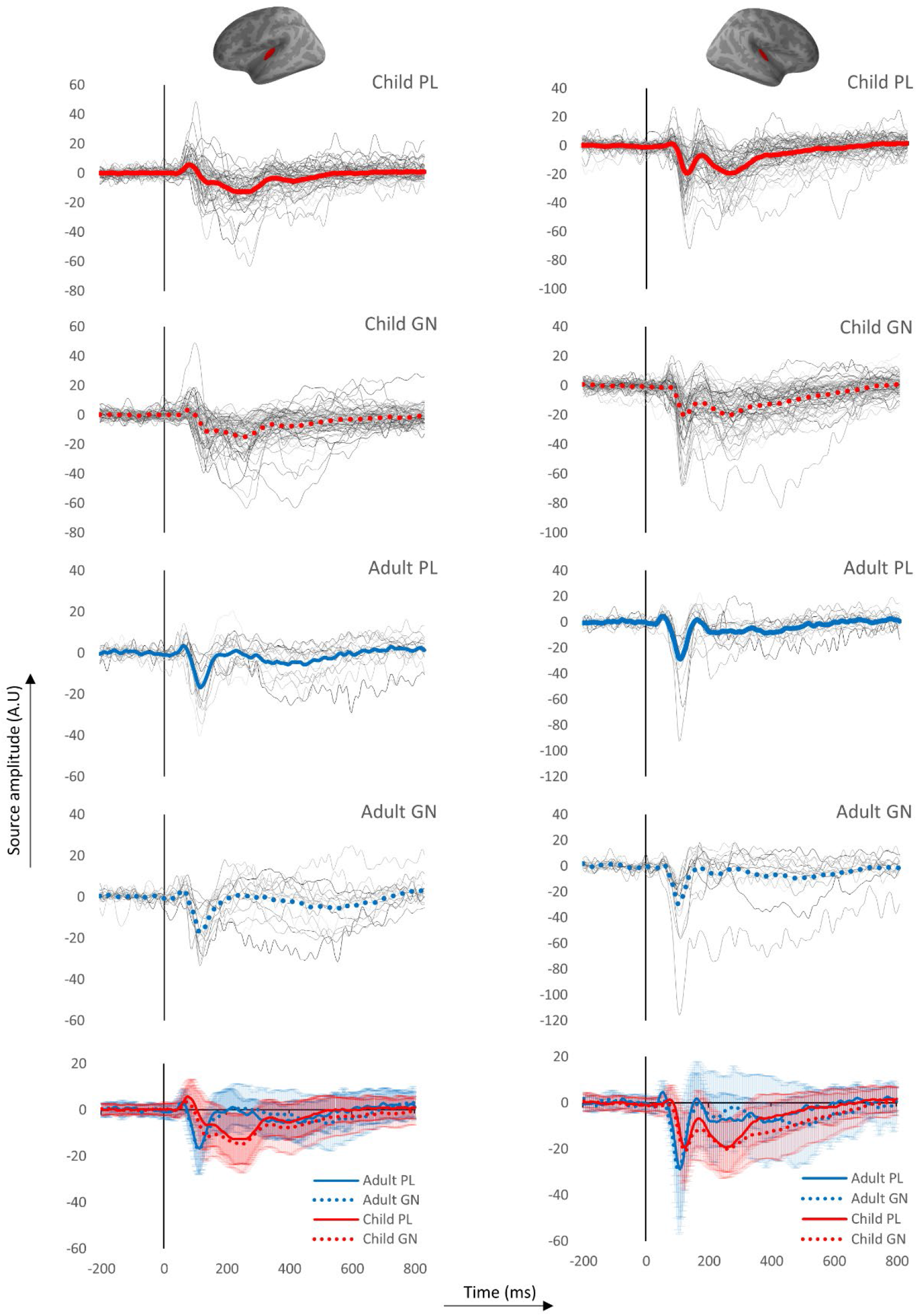
M/EEG combined Source estimates in the left and right transverse temporal gyrus (red area) of adults (blue lines) and children (red lines). Waveforms are an average of the entire area. Figures depict the passive (solid lines) and Inhibition (No-go) (dotted lines) waveforms in the left (top) and right (bottom) hemisphere. Shaded areas around the waveform represent the standard deviation (SD)

The activation pattern at ∼250ms in the auditory cortex is unique to the child brain (Fig. 3&4) in our data. Consequently, the activation pattern in children vs. adults in the 250-ms time-window reflects different brain regions, their strength is not directly comparable. Moreover, the activation pattern at ∼250ms in the auditory cortex is unique to the child brain (Fig. 3&4) in our data. Therefore, we did not directly contrast adults and children for this activation pattern. The behavioral relevance of this activation pattern in children is discussed elsewhere (van Bijnen et al., 2022). Due to the qualitatively different nature of the activation at 250 ms, the statistical analysis was limited to the earlier (50-200 ms) auditory cortex responses (P1-N1-P2)

### Passive vs Go/No-go

#### P1

In children both task (p < .001) and hemisphere (p = .001) significantly affected the amplitude of the P1. The task effect was similar in the two hemispheres as the task x hemisphere interaction was not significant (p = .301). More specifically, the P1 in the PL task was stronger compared to the P1 in GN task in the left (M = 7.69, SD = 8.13 vs. M = 4.38, SD = 7.86) and right hemisphere (M = 3.08, SD = 5.84 vs. M = 0.01, SD = 7.61). The P1 in children was significantly stronger in the left, compared to the right hemisphere in both the PL and GN task (p < .001). Finally, there was no main effect of group on the P1 (p = .296).

Adults showed the same task effect as children, but no hemisphere effect since the group x task interaction was not significant (p = .163) but the group x hemisphere interaction was significant (p = .012). The task effect was similar in the two hemispheres as the task x hemisphere interaction was not significant (p = .301)and the group x task x hemisphere interactions (p = .204) were not significant. Similar to the child group, the P1 in the PL task was stronger compared to the P1 in GN task in the left (M = 5.59, SD = 4.44 vs. M = 4.7, SD = 3.97) and right hemisphere (M = 6.13, SD = 4.87 vs. M = 2.91, SD = 4.02).

The P1 did not consistently correlate with age in children, with only the No-go P1 in the right-hemisphere reaching significance (r = -.28, p = .02). No significant correlations were found between the P1 in children and their behavioral performance. In adults however, stronger No-go P1 in the left hemisphere was related to a higher ICV (more variability) (r = .69, p = .003). The correlation matrix is presented in table 2 and the associated scatterplot in figure 6.

**Table 2.** Partial (correcting for age in children only) correlation between the P1 amplitude and behavioral measures. Significant correlations marked in bold.

|  | Child P1 |  |  | Adult P1 |  |  |
| --- | --- | --- | --- | --- | --- | --- |
|  | ICV | RA | SSRT | ICV | RA | SSRT |
| L PL | -.052 | .019 | -.010 | .117 | -.024 | -.116 |
| R PL | .146 | .180 | -.072 | .308 | .164 | -.047 |
| L GN | -.034 | -.005 | -.105 | **0.689\*\*** | .284 | .270 |
| R GN | .206 | .189 | -.020 | -.022 | -.162 | -.191 |

Note: ICV = intra-individual coefficient of variability, RA = response accuracy, SSRT = stop signal reaction time. \* $p < 0.05$ , \*\* $p < .01$ , \*\*\* $p < 0.001$ .

**Figure 5.**
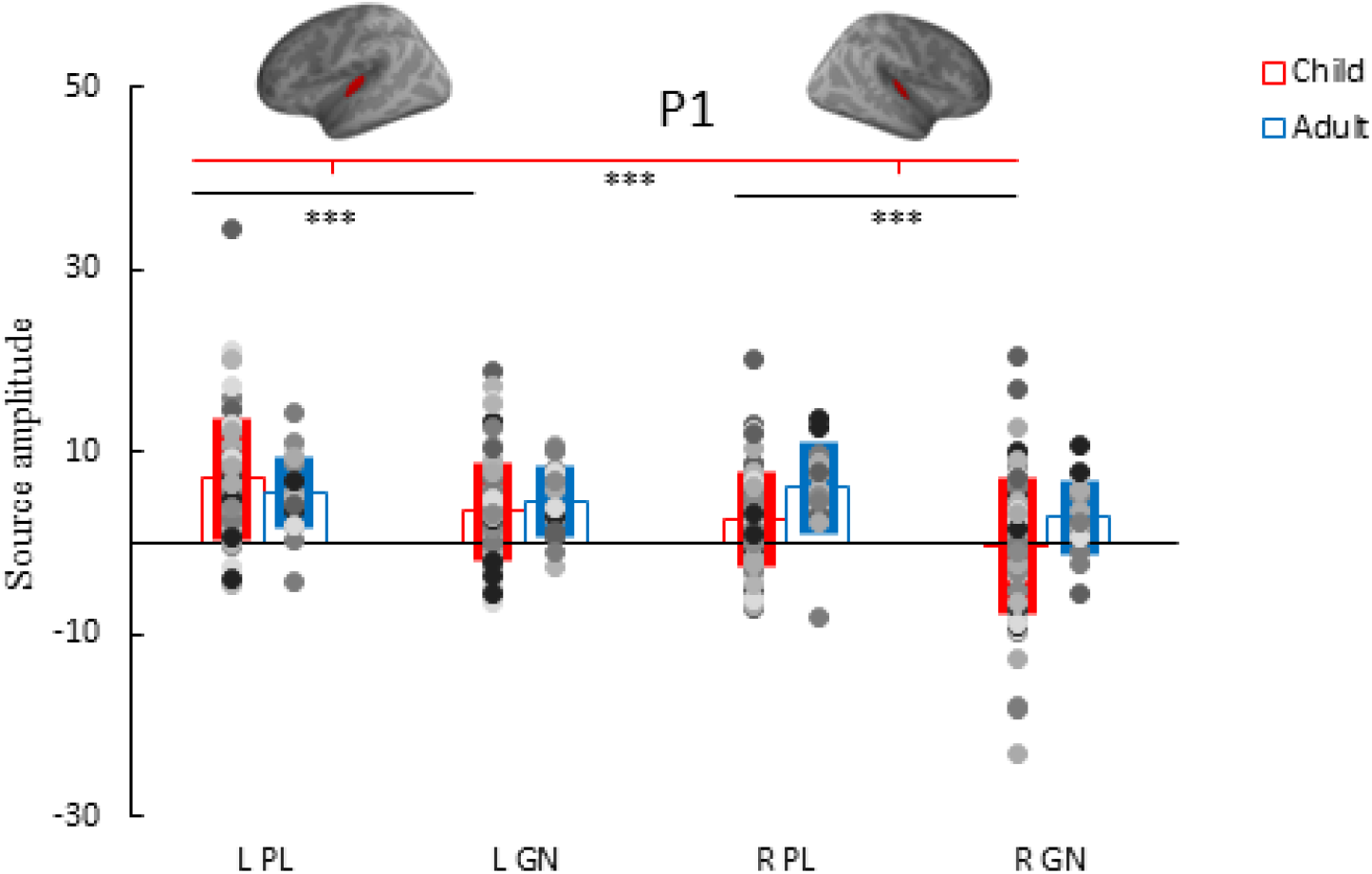
P1 amplitude individual data points, group average and standard deviation for the conditions: passive listening (PL) deviant tone and Go/No-go (GN) deviant tone in the left (L) and right (R) hemisphere of adults (blue) and children (red).

**Figure 6.**
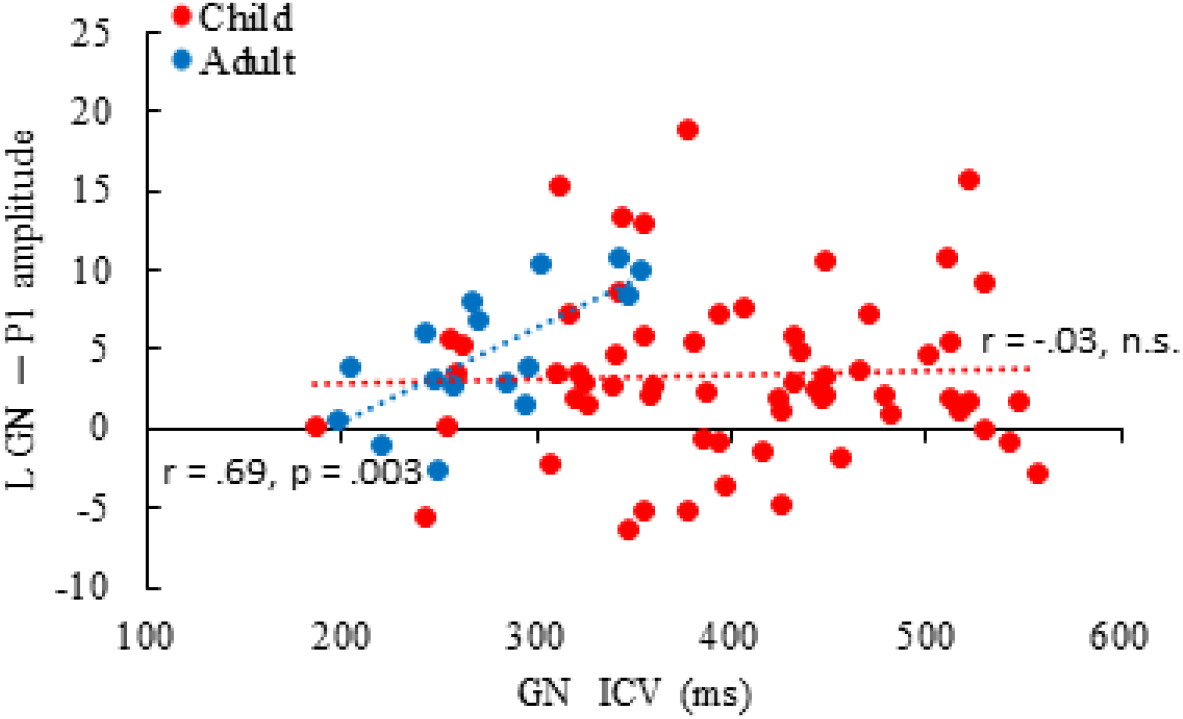
Scatterplot of the left-hemisphere P1 response to the No-go tone and the intraindividual coefficient of variability (ICV) of the Go/No-go (GN) task of children (red) and adults (blue)

#### N1

Children and adults showed similar N1 main effects as the N1 showed no significant interactions between group, task and hemisphere (p > .05). There were main effects of group and hemisphere, but the task did not influence N1 strength (p > .05). Adults showed stronger N1 responses compared to children (p = .044) and the right-hemisphere N1 responses were stronger compared to left-hemisphere N1 responses in both tasks and groups (p < .001) (Figure 7).

**Figure 7.**
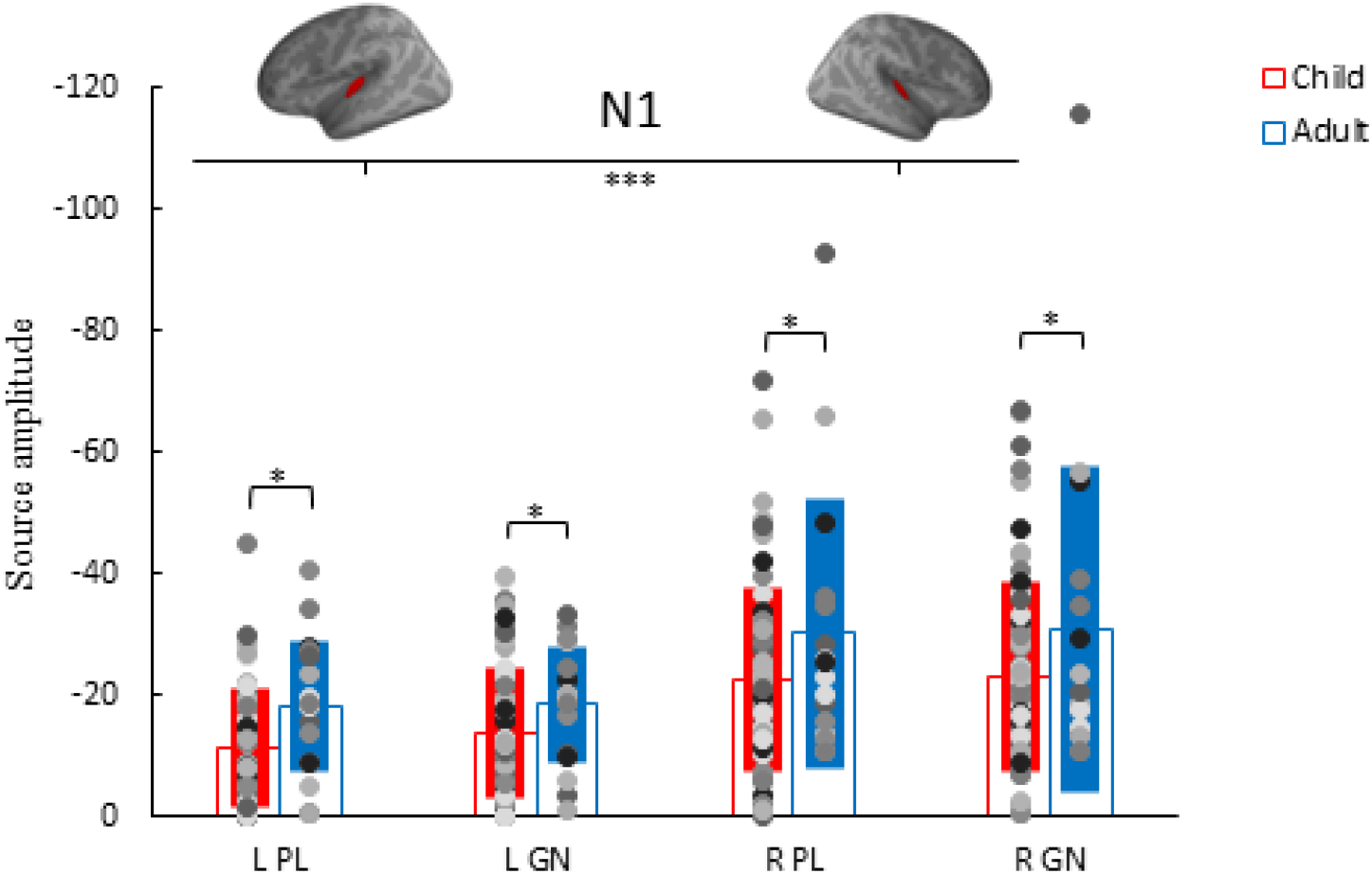
N1 amplitude individual data points (dots), group average (empty bar) and standard deviation (solid bar) for the conditions: passive listening (PL) deviant tone and Go/No-go (GN) deviant tone in the left (L) and right (R) hemisphere of adults (blue) and children (red).

The N1 did not consistently correlate with age in children, with only the No-go N1 in the right-hemisphere reaching significance (r = -.26, p = .035). In children, a stronger left-hemisphere N1 response in the PL task is associated with lower ICV (r = .302, p = .03). In contrast, adults show opposite associations with a stronger left-hemisphere N1 response associated with higher ICV (table 3) and poorer performance in both the PL (r = -.701, p = .003; r = -.67, p = .004, respectively) and GN (r = -.612, p = .012; r = -.793, p < .001, respectively) task. The correlation matrix is presented in table 3 and the associated scatterplots in figure 8.

**Table 3.** Partial (correcting for age in children only) correlation between the N1 amplitudes and behavioral measures. Significant correlations marked in bold.

|  | Child N1 |  |  | Adult N1 |  |  |
| --- | --- | --- | --- | --- | --- | --- |
|  | ICV | RA | SSRT | ICV | RA | SSRT |
| L PL | **.302\*** | .249 | .145 | **-.701\*\*** | **-.670\*\*** | -.394 |
| R PL | .082 | .101 | .092 | .067 | -.148 | .062 |
| L GN | .222 | .231 | .087 | **-.612\*** | **-.793\*\*\*** | -.448 |
| R GN | .102 | .107 | .108 | -.062 | -.290 | .017 |

Note: ICV = intra-individual coefficient of variability, RA = response accuracy, SSRT = stop signal reaction time. \* $p < 0.05$ , \*\* $p < .01$ , \*\*\* $p < 0.001$ .

**Figure 8.**
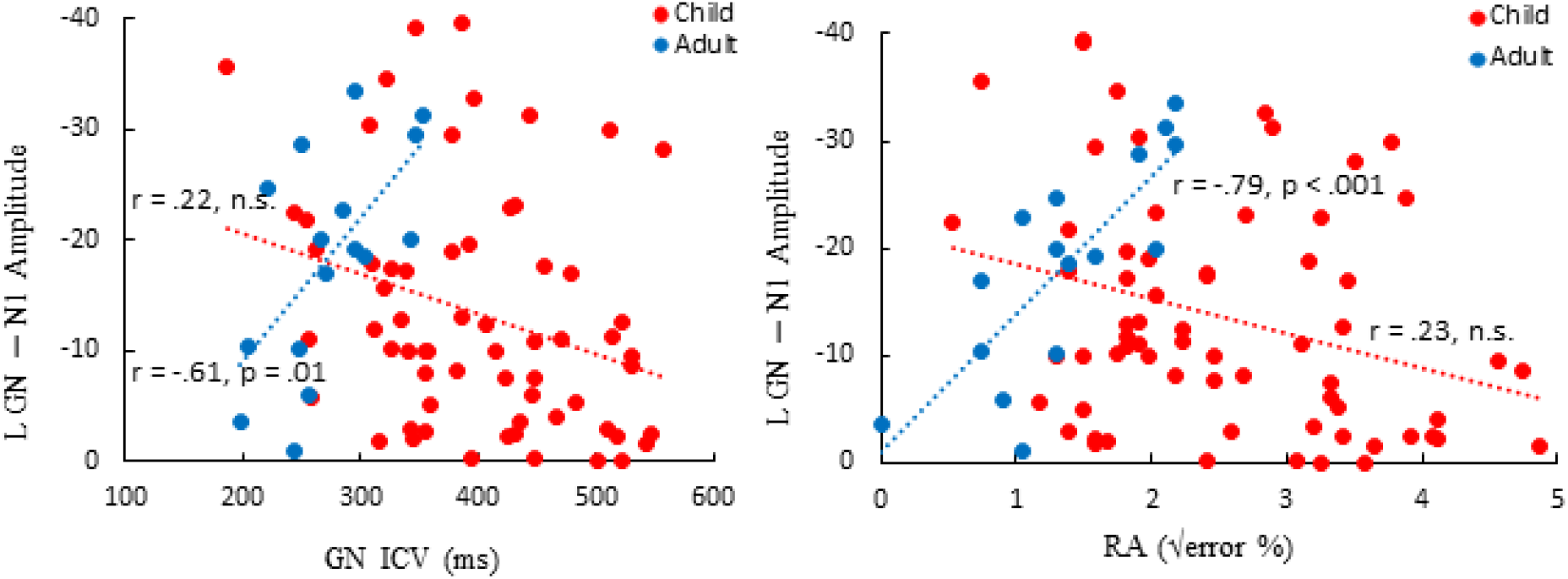
Scatterplot of the left-hemisphere N1 responses to the No-go tone and the behavioral performance measures of the Go/No-go (GN) task: intraindividual coefficient of variability (ICV; left) and response accuracy (RA; right) of children (red) and adults (blue)

#### P2

Similar to the N1, the P2 showed no significant interaction between group, task and hemisphere (p > .05). There were main effects of group and hemisphere, but task did not significantly effect P2 amplitudes (p > .05). Adults showed stronger P2 responses compared to children (p < .001) and the right-hemisphere P2 responses were stronger compared to left-hemisphere N1m responses in both tasks and groups (p < .001) (figure 9). The P2 showed consistent age effects with the correlation coefficient ranging between .3 and .351 (p ≤ 0.014).

**Figure 9.**
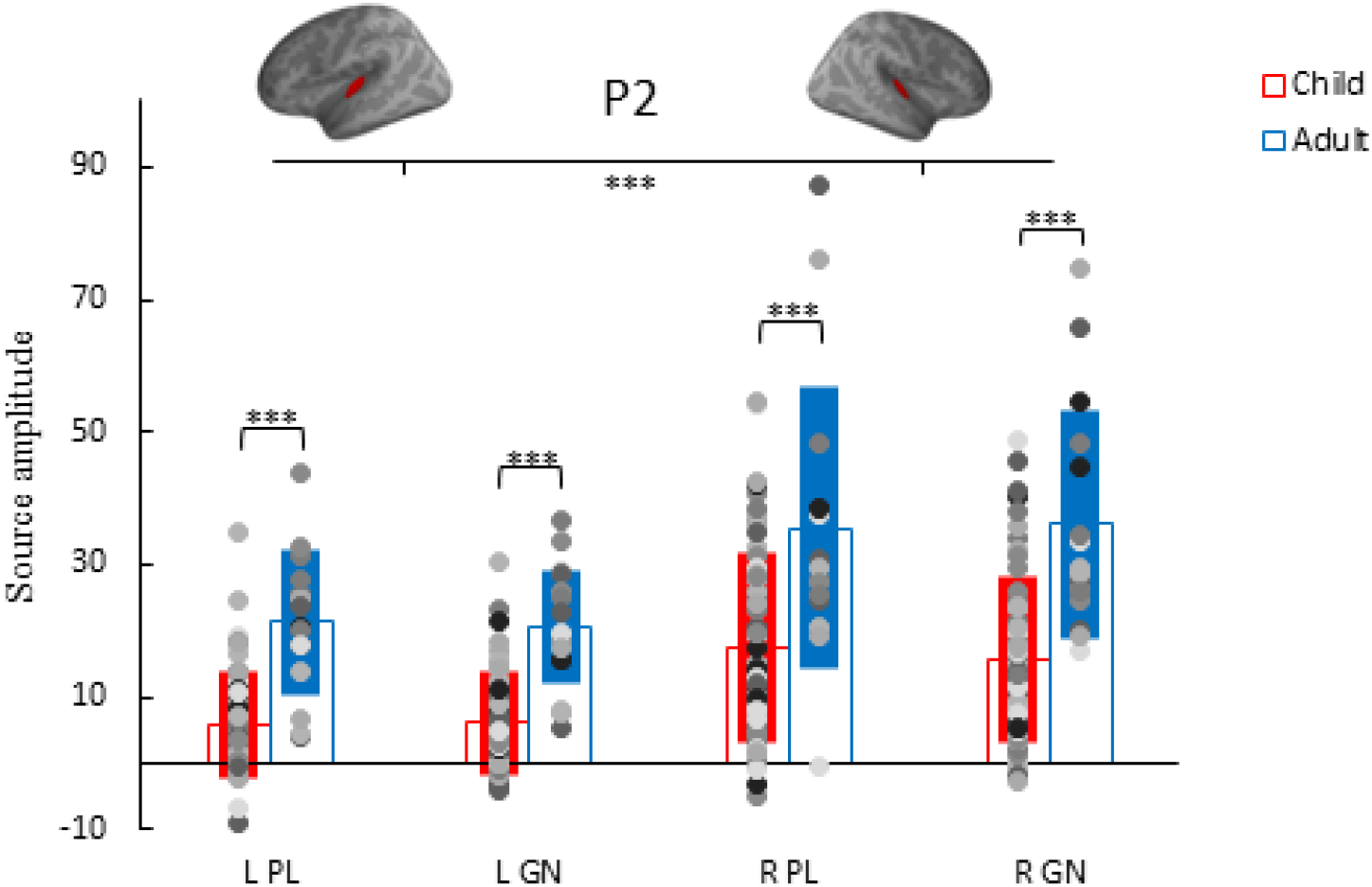
P2 amplitude individual data points (dots), group average (empty bar) and standard deviation (solid bar) for the conditions: passive listening (PL) deviant tone and Go/No-go (GN) deviant tone in the left (L) and right (R) hemisphere of adults (blue) and children (red).

The (partial) correlation revealed that, in children, a stronger No-go P2 in the right hemisphere is associated with lower SSRTs (r = -331, p = .008). In adults, left-hemisphere No-go P2 responses were associated with higher SSRT (r = .55, p = .027), higher ICV (r = .538, p = .031) and worse response accuracy (r = .535, p = .033). The correlation matrix is presented in table 4 and the associated scatterplots in figure 10.

**Table 4.**
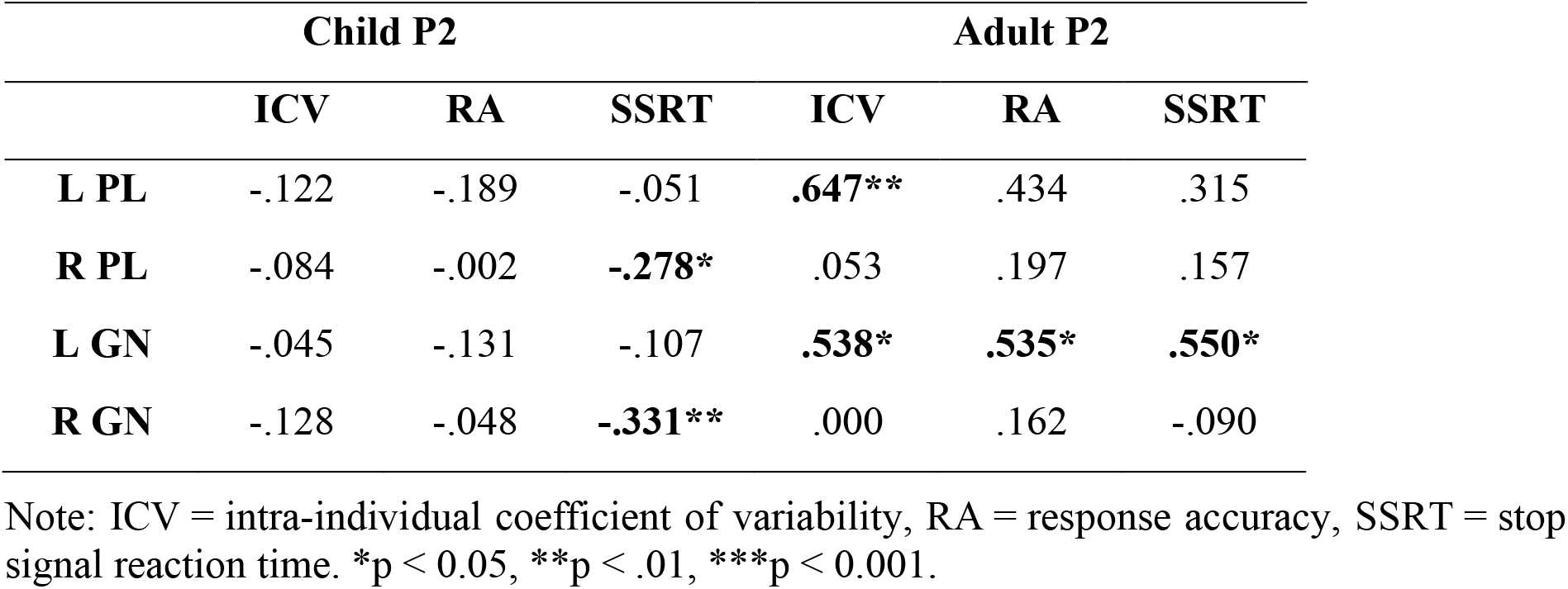
Partial (correcting for age in children only) correlation between the P2 amplitudes and behavioral measures. Significant correlations marked in bold.

**Figure 10.**
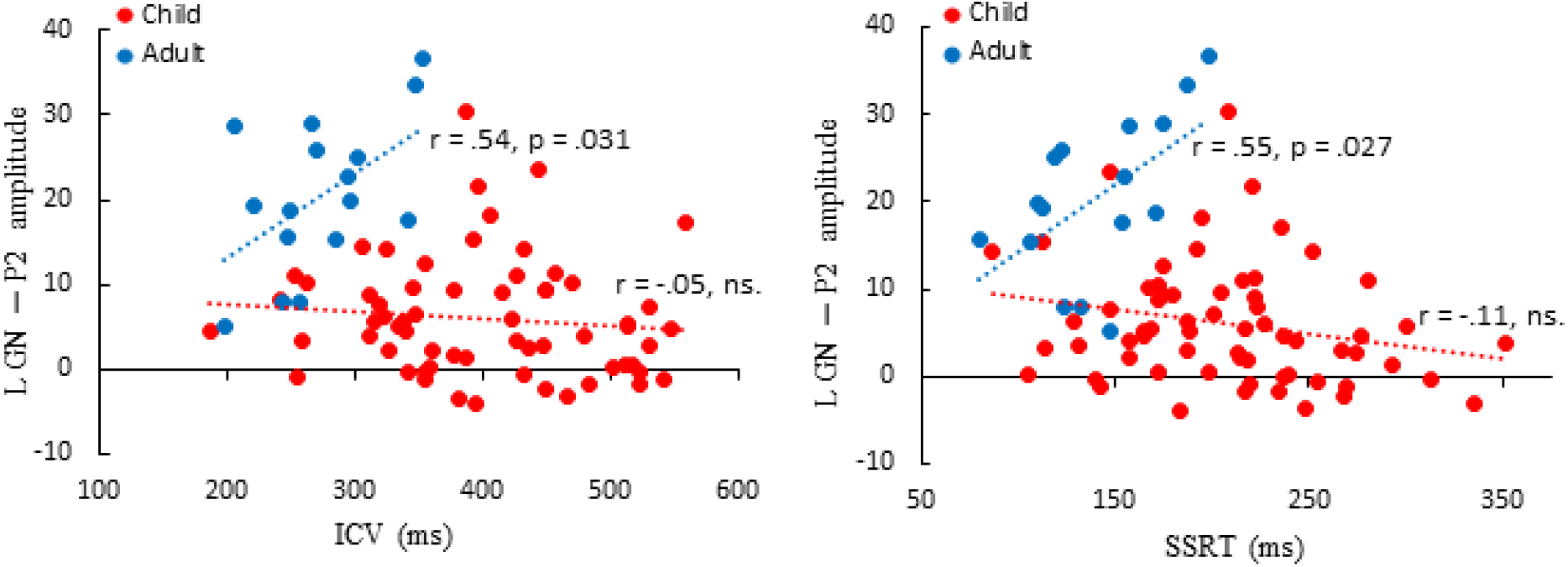
Scatterplot of the left-hemisphere P2 responses to the No-go tone and the behavioral performance measures of the Go/No-go (GN) task: intraindividual coefficient of variability (ICV; left) and stop-signal reaction times (SSRT; right) of children (red) and adults (blue)

## Discussion

In this study we found key differences in the auditory processing of adults and children. First, children rely on auditory areas for an extended period of time to process auditory information during a Go/No-go task compared to adults; the adult activation pattern shifts at ∼200ms from auditory to medial regions of the cerebral cortex that are implicated in cognitive control processing, whereas children show prolonged obligatory responses in the auditory cortex. This indicates a qualitative difference in cognitive control processing between adults and children. Second, the associations between the early auditory cortical responses and inhibition performance measures diverge between adults and children; while in children the brain-behavior associations are not significant, or stronger responses are beneficial, adults show negative associations between auditory cortical responses and inhibitory performance. Together this likely translates to a functional difference between adults and children in the cortical resources for performance consistency in auditory-based cognitive control tasks.

The developmental changes that have been indicated to reflect more efficient auditory processing are a more pronounced N1 and in general a gradual temporal dissociation of the earlier responses (P1-N1-P2) and an attenuation of the later N2 response (Ponton et al., 2000; Albrecht et al., 2000; Čeponienė et al., 2002; Takeshita et al., 2002; Wunderlich and Cone-Wesson, 2006; Sussman et al., 2008). The age-related dynamics, specifically the decrease in P1 at the early stages of development and a subsequent increase in N1, are in line with the present findings and are together likely to reflect maturational differences in the cortical circuitry (Orekhova et al., 2012). Notably, deep layers (lower III to IV) in the auditory cortex mature earlier, between 6 months and 5 years of age, compared to the superficial layers (upper III and II) that mature somewhere between 6─12 years (Ponton et al., 1999; Eggermont & Ponton, 2003; Moore & Guan, 2001; Moore & Linthicum, 2007). As we studied older children, their cortical P1 generators arguably closely reflect that of adults, unlike the neural generators of the N1 that are still developing in our sample of children. Accordingly, we found no age or group effects of the P1, but did for the N1 and P2. A similar explanation could account for our task effect in the P1 that showed smaller amplitudes to the No-go compared to the passive tones in both adults and children, but not in the N1 and P2. Indeed, highlighting the different neural generators of the P1 and N1/P2, a study modelling the adult auditory responses reported that an initial excitatory thalamocortical feedforward drive to layer II/III and V, via layer IV induced the P1. In contrast, a cortico-cortical feedback input to supragranular layers with a subsequent second feedforward input induced the N1 and P2 respectively (Kohl et al., 2019). Apparently, the necessity to inhibit a response lowers the thalamocortical input that is associated with the P1.

In addition to the amplitude and latency changes to the auditory responses, their function also changes; children seem to rely more strongly on sensory activation in active (inhibition) tasks. We show that, while the adult’s early auditory components (i.e., P1-N1-P2) are all negatively related to inhibitory performance, the early responses in children are not, or are (weakly) positively related. Instead, the child-specific auditory activation pattern at ∼250 ms post-stimulation (N2) in the left-hemisphere is reported to be strongly (positively) related to inhibitory performance (Johnstone et al., 1996; van Bijnen et al., 2022).

Earlier studies have highlighted the discrepancies between the early (P1-N1-P2) and the later, obligatory, (N2) response pattern in children, calling it “the additional process” (Johnstone et al., 1996) and suggesting it reflects the child’s wider range of attentional focus, resulting in similar neural response patterns in distinct situations (e.g., active vs passive) while still behaviorally relevant (Johnstone et al., 1996; van Bijnen et al., 2022). Our data supports this claim as we were unable to falsify that the early response amplitudes are not behaviorally relevant for inhibitory performance in children. Thus, the child-specific, obligatory, N2 response seems to be unique in its behavioral relevance compared to the earlier obligatory auditory responses in children. Another possibility is that the early auditory child responses are also beneficial for task performance, but perhaps the child and adult brain-behavior associations are diametrically opposed. This would complicate matters; as the brain-behavior associations possibly “flip” during development and this could limit the ability to detect brain-behavior associations in the P1-N1-P2 complex in children.

Our data are in line with previous studies on the adult neurobiological mechanisms of response inhibition and shows it is remarkably different in children in auditory based tasks. In adults, response inhibition is supported by a broad frontoparietal network (Weiss & Luciana, 2022; Puiu et al., 2020) and similarly the midcingulate cortex is reported to be a major neural generator of the adult N2 in active tasks (Huster et al., 2010). However, we show that, in children, the major source of activation in this time-window during No-go trials is in the auditory cortex. It suggests that this auditory activation pattern in children becomes obsolete as the brain becomes more efficient in discriminating auditory stimuli and determining their behavioral relevance. The child brain is not merely an “miniature” adult brain but the mechanisms that govern inhibitory performance in children are functionally distinct from adults. The maturational changes in the auditory response coincide with improvement in inhibitory performance during childhood and adolescence, but this transition is likely aided by child-unique mechanisms.

This study cannot give a causal explanation for the neural and behavioral changes during development. However, this is a comprehensive study that contrasted source models of adult and child auditory processing and investigated the relevance for response inhibition. The experiment was carefully designed to limit external factors (e.g., differences in motor and/or stimuli between conditions). Thus, this is the most direct comparison possible between active and passive tasks. In combination with our methods (combined M/EEG and individual MRIs) it provides greater confidence in our conclusion that associations between auditory activation and inhibition task performance differ in adults and children.

Future studies should look at individual changes over time (i.e., longitudinal) to investigate whether the developmental changes relate to the improved performance in response inhibition. Ideally, this should also include magnetic resonance spectroscopy (MRS) to measure the biochemical changes in the brain, as maturation of GABAergic vs. glutamatergic circuits likely play a crucial role in both auditory and response inhibition development (Le Magueresse & Monyer, 2013; Sanes & Kotak, 2011; Du et al., 2016; Silveri et al., 2013). Moreover, an important remaining question is whether the effects are limited to the left-hemisphere, or depend on handedness.

The clinical importance of achieving competent response inhibition highlights the value of a complete understanding of the typical development of this process. This study shows remarkable differences between adults and children in (auditory) processing during response inhibition. It emphasizes the cruciality of sensory processing during (critical) periods of development until adult-like response inhibition networks have matured sufficiently. Future studies that examine the source models longitudinally, and account for biochemical changes, would be welcomed.

## Acknowledgements

We are grateful to Hanna-Maija Lapinkero, Suvi Karjalainen, Maria Vesterinen & Janne Rajaniemi for help with data collection and to Amit Jaiswal, Erkka Heinilä and Jukka Nenonen for their help with preprocessing and scripting and Joona Muotka for help with the statistical analysis. This work was supported by EU project ChildBrain (Horizon2020 Marie Skłodowska-Curie Action (MSCA) Innovative Training Network (ITN) – European Training Network (ETN), grant agreement no. 641652) and the Academy of Finland grant number 311877.

